# Neuronal Activity-Dependent Electroosmosis and Its Potential Role in Interstitial Fluid Flow in Glymphatic System

**DOI:** 10.64898/2026.06.14.732157

**Authors:** Parnian Hemmati, Anthony C. Wang, Mayumi L. Prins, Christopher C. Giza, H. Pirouz Kavehpour

## Abstract

The mechanisms driving interstitial fluid flow through brain parenchyma remain unresolved, limiting our understanding of how fluid transport contributes to glymphatic waste clearance and broader aspects of brain metabolism and neuronal activity. Existing theories based on diffusion or pressure gradients fail to explain sustained, directional flow through the tortuous extracellular space (ECS), particularly under normal physiologic conditions. Here we propose that electroosmosis, fluid motion driven by endogenous electric fields acting on charged brain tissue, provides a biophysically consistent mechanism for intraparenchymal interstitial fluid transport while generating pressure distributions favorable for periarterial influx and perivenous efflux. Using computational modeling informed by anatomical reconstructions of ECS microstructure and local field potential (LFP) recordings, we show that electroosmotic flow generates physiologically realistic velocities and reproduces brain state dependent differences in glymphatic transport, including the enhanced glymphatic flow observed during sleep compared to wakefulness. A physics-informed reduced-order model (ROM) further demonstrates that these microstructure-resolved results upscale consistently to tissue-level transport. Moreover, electroosmotic flow can induce directional pressure gradients across perivascular interfaces, facilitating both influx and efflux. This mechanism provides a unifying framework linking neuronal activity, parenchymal flow, and compartmental pressure regulation. In contrast, pressure gradients substantially larger than physiological estimates generated much smaller velocities and failed to account for the observed transport rates. These findings address a major gap in glymphatic physiology and suggest that modulation of electric field properties, via endogenous activity or external neuromodulation, could serve as a therapeutic strategy to enhance solute clearance in neurological disorders.

**Significance Statement:** A major challenge in brain physiology is the lack of a unifying physical model that explains how fluid moves through the narrow and tortuous extracellular space; a process essential for nutrient distribution and waste clearance. Existing frameworks cannot account for sustained, directional transport under normal physiological conditions. We show that electroosmosis (fluid motion generated when endogenous neuronal electric fields act on charged cellular surfaces) provides a biophysically consistent mechanism for this transport. Using realistic extracellular microstructures and a validated tissue scale reduced order model, our model reproduces experimentally observed brain state dependent transport. Establishing electroosmosis as a possible contributor to interstitial fluid movement offers a new conceptual basis for linking neuronal activity to fluid circulation and for guiding strategies to enhance brain clearance.

## Introduction

The glymphatic system is a brain-wide clearance pathway that transports cerebrospinal fluid (CSF) through the central nervous system, facilitating solute exchange and removal of neurotoxic waste products ^1–3^ (Fig. 1a). Dysfunction of this system has been implicated in neurological disorders ranging from Alzheimer’s disease to traumatic brain injury, where impaired clearance accelerates neurodegeneration ^4–7^. Understanding the mechanisms that regulate interstitial fluid (ISF) transport, particularly within the brain parenchyma, is therefore essential both for advancing basic physiology and for identifying therapeutic strategies that enhance waste removal. The glymphatic system is commonly divided into three segments: influx of CSF along periarterial spaces, transport through the parenchymal extracellular space (ECS), and efflux along perivenous pathways ^11;12^. While arterial pulsations and perivascular pressure gradients have been proposed as drivers of periarterial influx ^13^, the mechanism sustaining parenchymal flow remains unresolved. An important distinction in glymphatic physiology is that transport within the brain parenchyma should not necessarily be viewed as a direct continuation of fluid circulation within perivascular spaces (Fig. 1e). Periarterial and perivenous pathways form relatively open conduits surrounding blood vessels, where arterial pulsations, vasomotion, and other vascular oscillations can effectively drive fluid motion. In contrast, the parenchymal extracellular space consists of a highly confined and tortuous network of nanoscale channels whose hydraulic resistance is substantially greater than that of perivascular pathways. Consequently, only a fraction of the fluid circulating within perivascular spaces is expected to exchange with the parenchyma at any given location. The physical mechanisms governing transport within these two compartments may therefore be fundamentally different. While vascular pulsations may contribute to circulation within perivascular pathways and facilitate fluid exchange at tissue boundaries, the mechanism responsible for sustaining directional transport through the highly resistive extracellular space remains unresolved. The ECS (Fig. 1b), is a narrow, tortuous microenvironment, with widths on the order of tens to hundreds of nanometers, posing significant resistance to fluid motion ^14;15^.

**Figure 1:**
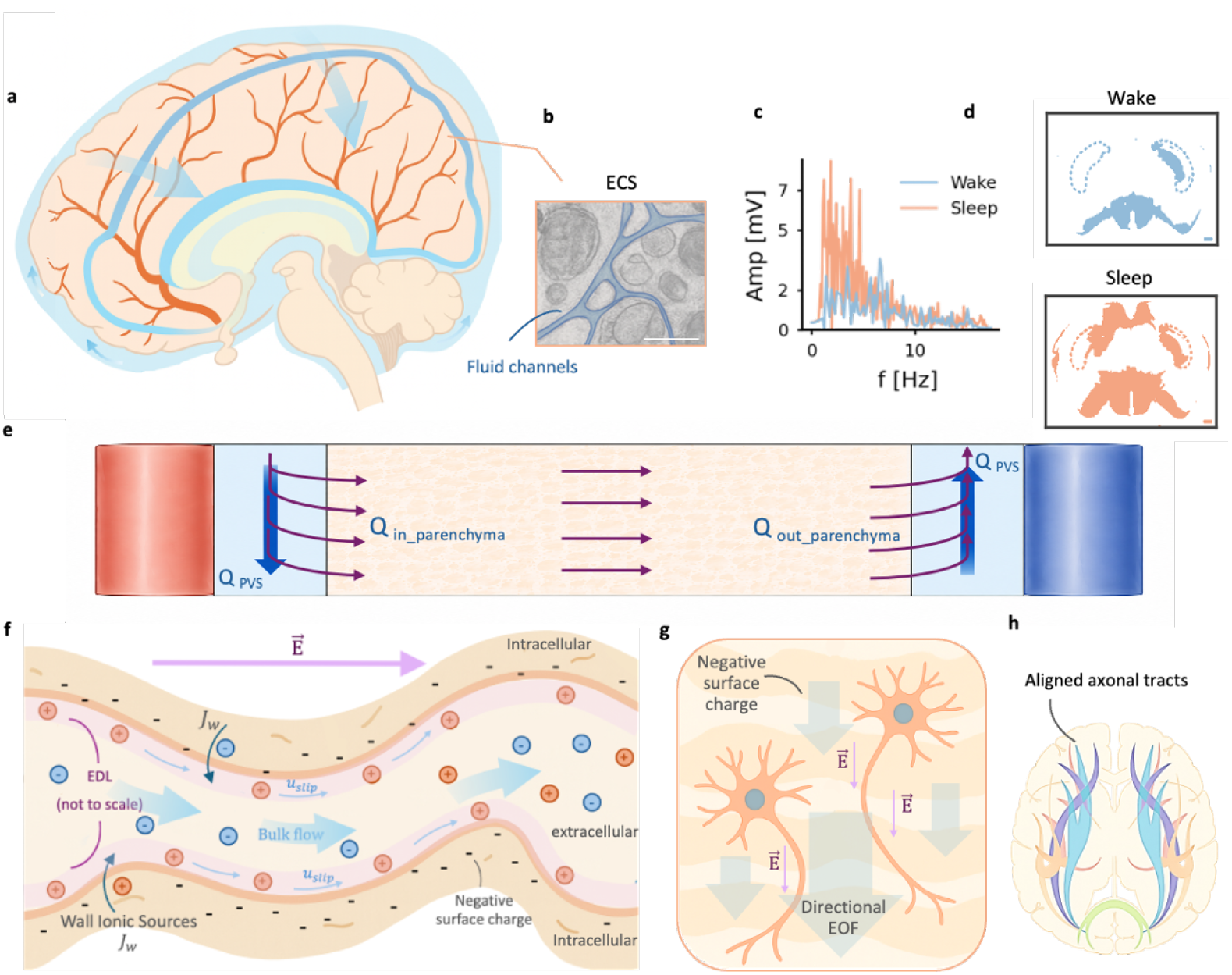
Electroosmotic hypothesis of glymphatic transport. **a**. Schematic of glymphatic circulation in human brain. **b**. EM of cryo-fixed brain tissue from adult mouse cerebral cortex. The ECS is pseudo-coloured blue. Scale bar 1 µm. adapted from ^8^ **c**. Power spectra of local field potentials during wake and sleep, demonstrating a shift toward higher-amplitude, lower-frequency oscillations in sleep. Data taken from ^9^. **d**. State-dependent tracer penetration into parenchyma, with deeper glymphatic transport during sleep compared to wake ^9^. **e**. Conceptual distinction between perivascular and parenchymal transport. Fluid circulates within periarterial and perivenous spaces, while only a fraction exchanges with the ECS. **f**. Schematic of electroosmotic flow: electric fields act on positive charged ions lining ECS walls, producing slip velocity (*u*_slip_) that drives near plug-like bulk flow. **g**. Neuronal activity generates spatially varying electric fields that interact with charged surfaces to create directional EO flow. **h**. Structural anisotropy of aligned axonal tracts ^10^ guides field propagation, channeling flow along preferred tissue orientations.

Classical studies established that diffusion in the ECS is strongly influenced by geometric factors such as tortuosity and volume fraction, which hinder the movement of macromolecules relative to free solution ^16;17^. More recent high-resolution imaging has revealed that the ECS forms a heterogeneous nanoscale network whose structure varies dynamically across physiological states ^18;19^. Ultrastructural analyses have also demonstrated that traditional chemical fixation methods can significantly underestimate ECS volume, whereas cryo-fixation preserves a larger and more physiologically representative extracellular space ^8^. Together, these studies highlight the importance of realistic ECS geometry when modeling interstitial transport in brain tissue.

Pressure-driven bulk flow has been proposed as a mechanism for intraparenchymal transport, but there is little direct evidence supporting the presence of persistent pressure gradients capable of generating physiologically relevant flow through parenchyma. Moreover, recent computational reconstructions argue against pressure-driven intraparenchymal flow. Holter *et al*. ^20^ simulated interstitial flow in 3D electron microscopy–based ECS geometries and found that even under unrealistically large pressure gradients predicted flow velocities were, several orders of magnitude below tracer penetration rates observed in vivo. These results demonstrate that pressure gradients within the parenchyma are too small, and tissue permeability too low, to sustain meaningful bulk flow. Diffusion, although always present, is nondirectional and too slow to account for the observed rates of solute transport. Specifically, diffusion through reconstructed ECS geometries can account for gradual solute spread over hundreds of micrometers, but only at timescales of tens of minutes to hours ^20^. While this may suffice for small ions, it is far too slow and nondirectional to account for the rapid, state-dependent tracer transport observed experimentally ^3;9;21^. Thus, diffusion alone cannot provide the directional and dynamic regulation of glymphatic flow that is required in vivo. Likewise, persistent pressure gradients sufficient to drive bulk parenchymal flow have not been directly observed. Neuronal activity can also induce osmotic shifts and transient cell swelling due to ion redistribution across cellular membranes, altering ECS volume fraction ^22^. However, such processes primarily redistribute water across membranes and do not inherently generate sustained directional flow within the ECS. As a result, the fundamental driver of intraparenchymal transport remains a major gap in glymphatic physiology.

In general, waste accumulation within the parenchyma increases local osmotic pressure relative to incoming cerebrospinal fluid, creating a tendency for water to move into, and remain within, the tissue. While such osmotic forces may facilitate fluid entry from periarterial spaces, they do not inherently promote clearance toward downstream efflux pathways and may, in some circumstances, oppose it. Consequently, efficient glymphatic transport requires not only a mechanism capable of driving directional interstitial flow but also one capable of overcoming or balancing the osmotic and hydraulic constraints associated with continuous solute clearance. These considerations further suggest that passive pressure- and diffusion-based mechanisms alone may be insufficient to fully explain the observed efficiency and directionality of glymphatic transport.

Recent experimental observations suggest a link between neuronal dynamics and ISF movement. Imaging studies show that glymphatic transport is enhanced during sleep compared to wakefulness ^21;23^, coinciding with shifts in neuronal activity toward high-amplitude, low-frequency oscillations. In particular, Jiang-Xie *et al*. ^9^ demonstrated that state-dependent changes in local field potentials (LFPs) directly correlate with CSF penetration, implicating electrical activity as a modulator of glymphatic function. Recorded LFPs during non-rapid eye movement (NREM) sleep have higher amplitudes and dominant low frequency compared to wake, as shown in Fig. 1c. Correspondingly, CSF penetration into the parenchyma is markedly increased during NREM (Fig. 1d). Moreover, local neuronal silencing reduced CSF influx in the same brain state, isolating neuronal activity as a causal driver of fluid movement. Recent human studies further support a link between neuronal activity and glymphatic function, showing that sleep-associated increases in EEG delta power and reductions in beta power are accompanied by enhanced glymphatic transport and reduced parenchymal resistance ^24^. Together, these findings support the idea that endogenous brain electric fields play a causal role in parenchymal fluid dynamics.

Here we propose that electroosmosis (EO), the motion of fluid driven by electric fields acting on charged surfaces, provides a biophysically consistent mechanism for ISF transport through the ECS. EO is not an unfamiliar phenomenon in physiology; it has been observed in other biological tissues where charged porous matrices interact with endogenous electric fields, establishing its plausibility as a transport mechanism ^25–27^. The negatively charged extracellular matrix naturally satisfies the conditions for EO ^28^, while ongoing neuronal activity generates time-varying electric fields capable of driving fluid motion ^29–31^. Using anatomically reconstructed ECS geometries ^32^ and computational multiphysics simulations informed by experimental LFP recordings, we show that electroosmotic flow produces physiologically realistic velocities, reproduces brain state dependent differences in intraparenchymal ISF transport, and intrinsically generates pressure gradients that reinforce influx and efflux. A physics informed reduced order model further demonstrates that these results extend consistently to tissue-scale transport. To our knowledge, this is the first quantitative study investigating electroosmosis as a contributor to glymphatic transport through brain parenchyma. Beyond its physiological relevance, an EO-based mechanism for parenchymal transport has broad implications for translational neuroscience: by linking neuronal activity to fluid dynamics, it suggests strategies to modulate glymphatic function for therapeutic benefit, with potential applications in enhancing drug delivery and promoting clearance of pathological solutes in neurodegenerative diseases ^13;33–35^.

## Results and Discussion

EO arises when an electric field acts on a charged porous medium, driving near plug-like fluid flow through confined channels. The interstitial space (ISS) of brain parenchyma naturally satisfies the prerequisites for this phenomenon: the extracellular matrix carries a net negative surface charge ^28^, and neuronal activity generates time-varying local electric fields ^29;30^.

A key feature of EO is the formation of an electric double layer (EDL) at charged surfaces. The characteristic EDL thickness, the Debye length, is given by 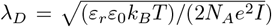, where *ε*_*r*_ is the relative permittivity, *ε*_0_ the vacuum permittivity, *k*_*B*_ Boltzmann’s constant, *T* the temperature, *N*_*A*_ Avogadro’s number, *e* the electron charge, and *I* the ionic strength. For physiologic ionic concentrations, *λ*_*D*_ ≈ 1 nm, much smaller than the typical ECS channel widths (10–100 nm). This places the brain firmly in the thin-EDL regime, where EO can be modeled by an effective slip velocity at the wall rather than explicit ion transport. At negatively charged surfaces, counterions from the surrounding electrolyte accumulate within the electric double layer (EDL). When an electric field is applied tangentially along the wall, these counterions migrate downstream, entraining solvent molecules and generating a slip velocity at the interface. This interfacial slip, transmitted by viscous shear into the channel interior, produces the characteristic near plug-like velocity profile of electroosmotic flow (Fig. 1f). The Helmholtz–Smoluchowski relation formalizes this boundary condition as 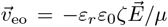 where *ζ* is the wall zeta potential, µ the fluid viscosity, and 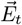 the tangential component of the electric field. This effective slip replaces the classical no-slip condition, providing the basis for intraparenchymal EO transport.

The electric fields in the brain are inherently dynamic, with magnitudes and frequencies governed by neuronal activity ^29^. To capture their effect on EO, we modeled the effective velocity spectrum as 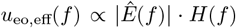 where 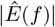 is the spectral magnitude of the field and *H*(*f*) is a low-pass transfer function representing frequency-dependent attenuation due to electric double layer relaxation and tissue-level low-pass filtering in the ECS ^36^. This framework predicts that low-frequency oscillations more efficiently drive EO, while higher-frequency components are suppressed. As shown in Fig. 1g, spatially varying electric fields generated by action potentials and oscillatory activity interact with nearby charged surfaces, aligning flow along the orientation of axonal tracts. This coupling results in local directionality of flow, ensuring that EO does not simply produce random pore scale motion but instead channels fluid preferentially along neuronal structures.

At a larger scale, the structural anisotropy of white matter bundles and aligned fiber tracts further organizes these flows (Fig. 1h) ^37^. By guiding field propagation across extended tissue domains, this anisotropy channels EO-driven transport from periarterial influx zones into deeper parenchyma and toward perivenous efflux routes. Together, these features integrate molecular physics, neuronal activity, and tissue architecture into a coherent framework for directional fluid movement.

Reported experimental results are consistent with this model: during sleep, LFPs show increased amplitude and dominant power at lower frequencies compared to wake (Fig. 1c). These state-dependent changes coincide with deeper CSF penetration into parenchyma (Fig. 1d), supporting the hypothesis that neuronal activity modulates glymphatic transport through EO.

To assess the feasibility of electroosmosis as a mechanism for parenchymal flow, we constructed a three-dimensional computational model of a 4 µm^3^ brain tissue block. The geometry was derived from electron microscopy reconstructions of hippocampal neuropil reported by Kinney *et al*. ^32^. As shown in Fig. 2a, a representative subvolume was cropped, surface-smoothed, and remeshed to improve quality for computational simulations. Since our focus was on fluid transport, the solid tissue domain was inverted to extract the fluid-accessible interstitial space, which was subsequently meshed (mesh statistics and independence analysis are provided in the Methods). The numerical implementation was independently validated in a microchannel system with experimental data reported by Peng *et al*. ^38^, validation results provided in Supplementary Fig. S1.

**Figure 2:**
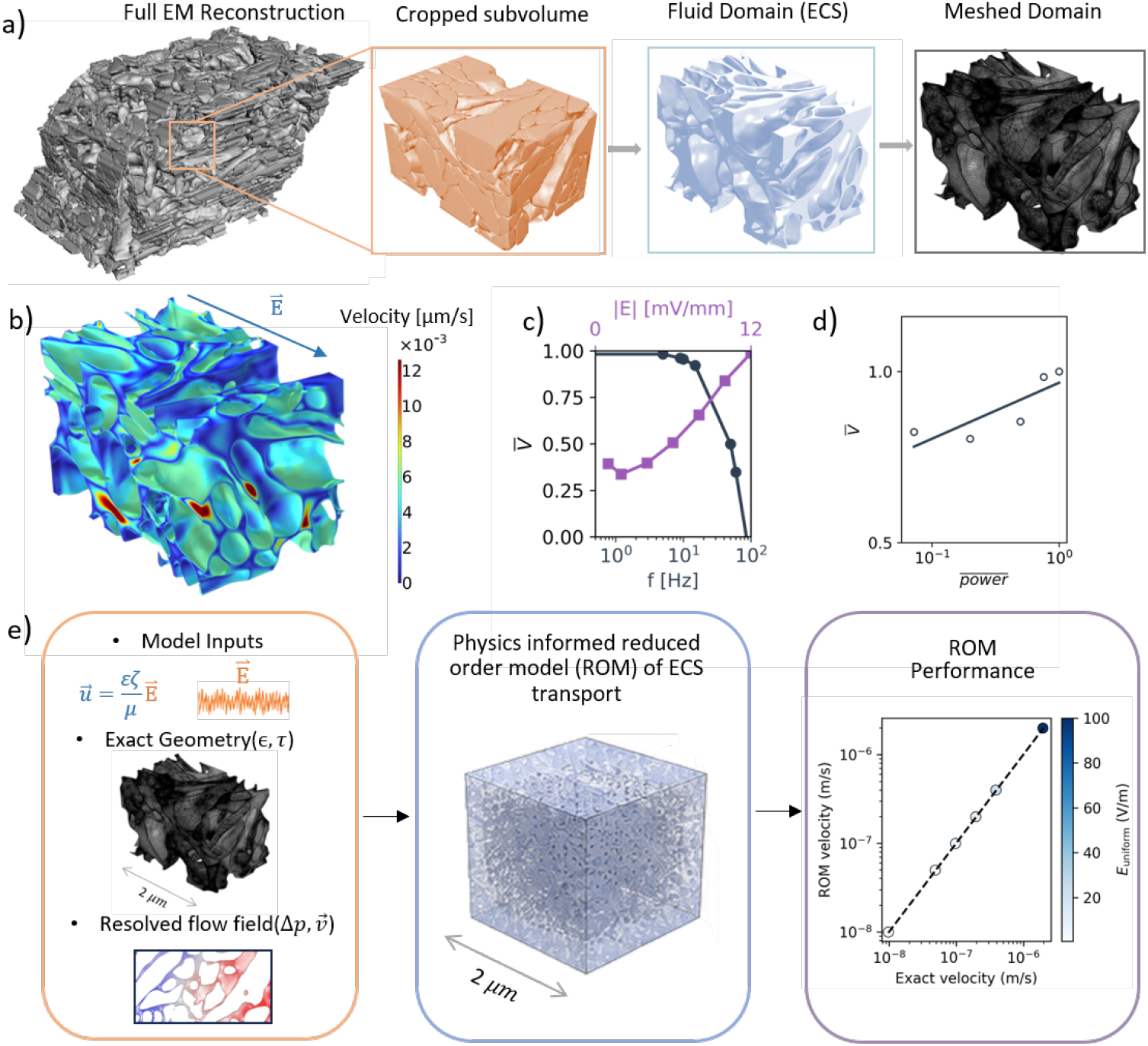
Electroosmotic flow in brain parenchymal microstructure. **a**, Workflow for geometry preparation: full EM reconstruction of hippocampal neuropil ^32^, cropped subvolume, extraction of fluid-accessible domain, and meshed interstitial space. **b**. Velocity field induced by electroosmosis within the meshed subvolume under an applied electric field. **c**. Parametric study of normalized electroosmotic velocity as a function of electric field frequency (*f*, Hz) and magnitude (|*E*|, mV/mm). 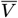 denotes velocity normalized by its maximum value across all tested conditions. **d**. Normalized average velocity increases with electric field power. 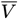 denotes velocity normalized by the maximum value reached by maximum power **e**. Physics informed ROM of ECS transport. Left panel: model inputs governing physics, Electric field (E), ECS geometry characterized by porosity and tortuosity, and the resolved microscale flow field. Center panel: the ROM operates on a homogenized 2 µm domain that can be upscaled to tissue dimensions. Right panel (ROM performance): predicted average transport velocity versus exact microstructure resolved velocity

To simulate electroosmotic flow within this geometry, we applied a zeta potential of −25 mV to the channel walls, consistent with reported brain’s extracellular wall potentials ^39^. (Sensitivity analyses of key electrokinetic parameters in model are reported in the Supplementary Information (Fig. S2). Symmetry boundary conditions were imposed on the four lateral faces to mimic unbounded tissue, while flow inlet and outlet conditions were applied along the direction of the externally applied electric field derived from local field potentials. (Model robustness under different boundary condition is studied and reported in the Supplementary Table S1.) The resulting electroosmotic flow field within the tissue block is shown in Fig. 2b, where fluid motion aligns with the applied electric field despite the tortuous pore structure. A simple scaling check using the Helmholtz–Smoluchowski relation with *ζ* = −25 mV, *ε* ≈ 80*ε*_0_, and *µ* = 10^−3^ Pa·s gives *u*_slip_ ∼ *εζE*/*µ*. This yields *u*_slip_ ≈ 0.002–0.018 *µ*m/s for *E* = 0.1–1 V/m, consistent with the thin-EDL expectation that EO scales linearly with *E* and *ζ* and is independent of channel size. The simulated average velocities fall within the scale required to account for tracer penetration rates observed in vivo. Although direct measurements of fluid velocity within parenchymal ECS are lacking, tracer studies imply transport on the order of micrometers per second based on penetration depths achieved over experimental timescales ^9;21^. These results support electroosmosis as a viable mechanism for interstitial flow. Fig. 2c shows streamlines along a channel in the direction of applied electric field. The insets show how EO-induced slip velocities at walls extend into the channel center, generating plug-like velocity profiles. Conservation of mass dictates that as the cross-sectional area of the channels increases, velocity decreases, producing spatial variability in flow magnitude while preserving net directional transport.

A parametric study (Fig. 2c) demonstrates that electroosmotic velocity increases with electric field magnitude but decreases with increasing field frequency. This reduction at higher frequencies reflects limited double-layer charging and field attenuation, constraining EO efficiency at physiologically relevant high signal frequencies. Electroosmotic flow depends on both the amplitude and frequency of the driving electric field. To quantify this, we used the power of a waveform as the time-averaged squared field magnitude: 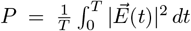. This metric integrates contributions from both field strength and oscillation frequency, providing a physiologically interpretable measure of neuronal activity. We generated synthetic families of sinusoidal waveforms spanning physiologic ranges of amplitude and frequency, guided by spectral features of recorded LFPs. When applied in simulations, increasing wave power led to proportionally higher EO velocities (Fig. 2d). These results establish wave power as a unifying parameter linking neuronal oscillations to EO-driven transport capacity. Microstructure resolved simulations, while mechanistically precise, are computationally prohibitive at tissue scales. To bridge this gap, we developed a physics-informed ROM of ECS transport (Fig. 2e). The ROM is trained through inputs including the governing EO equation, electric field, the ECS geometry characterized by porosity and tortuosity, and the resolved microscale flow field. As shown in the ROM performance panel (Fig. 2e, right), predicted average transport velocities agree closely with exact microstructure resolved velocities across a range of tested electric field magnitudes and brain states. Leveraging this, the calibrated ROM was applied to a tissue-scale domain of 200 *µ*m, constructed by tiling the unit 2 *µ*m ROM block (Fig. 3a). This large scale ROM construction, computationally intractable with the full microstructure resolved model, is what enables the brain state comparisons presented below. This ROM enables tissue scale simulations that retain the electrokinetic physics of the full model while reducing computational cost by several orders of magnitude.

**Figure 3:**
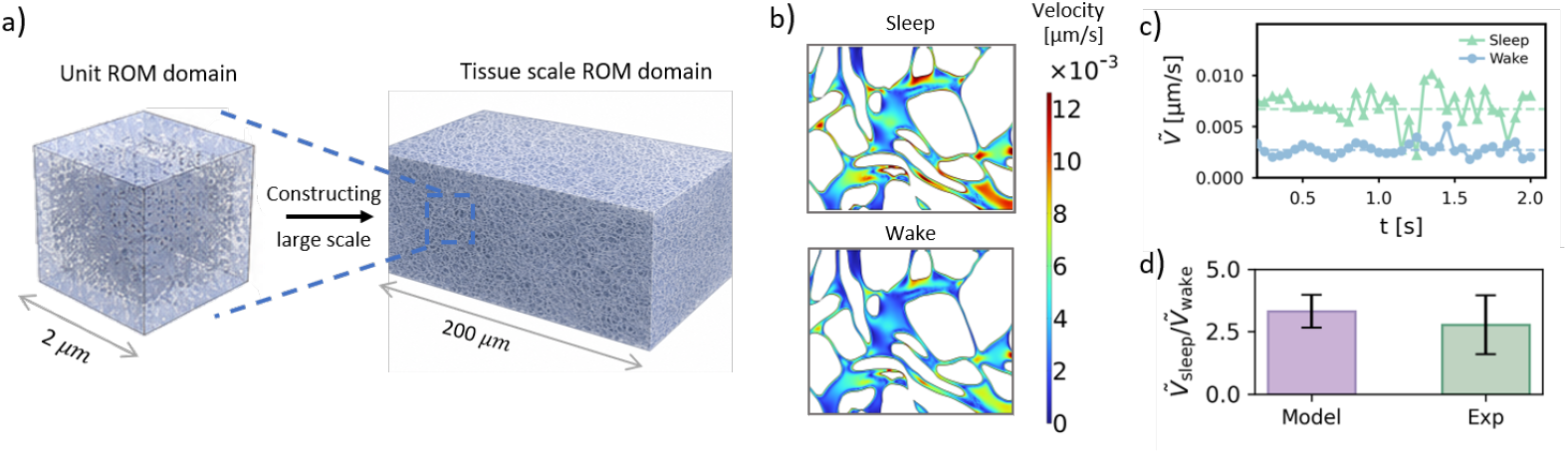
Brain-state dependence of electroosmotic flow. **a**. microstructure-resolved 2 *µ*m domain is homogenized into a tissue-scale ROM domain of 200 *µ*m. **b**. Velocity fields in the interstitial cross-sections under sleep and wake LFP inputs, showing higher velocities during sleep. **c**. Time dependent average velocity magnitude driven by oscillatory LFPs. 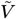 is the average velocity within the volume. **d**. Ratio of sleep-to-wake velocity from model compared with experimental data from Jiang-Xie et al. ^9^.

A critical prediction of the EO hypothesis is that glymphatic parenchymal flow should depend on neuronal brain state. In vivo imaging shows that intraparenchymal ISF flow is enhanced during sleep compared to wakefulness ^9;21^, coinciding with a shift in LFPs toward higher amplitude, lower frequency oscillations during sleep. To test whether EO can explain this correlation, we extracted LFP time-series from Jiang-Xie *et al*. and applied them as driving fields in our electroosmotic simulations of anatomically realistic ECS geometries ^32^.

The resulting velocity fields (Fig. 3b) show noticeably higher velocity magnitudes during sleep compared to wake. Physiological neural activity exhibits asymmetric, burst-like dynamics across multiple time scales, leading to nonzero effective transport over time. In addition, the highly heterogeneous and tortuous geometry of the extracellular space breaks spatial symmetry, allowing oscillatory electroosmotic forcing to generate net transport through rectification effects ^40^. As a result, time-dependent electroosmotic forcing can produce physiologically meaningful transport within realistic ECS geometries. Time-resolved velocities fluctuate with the oscillatory LFP input (Fig. 3c), but the time-averaged velocity during sleep is more than twice that during wake. Comparison with experimental tracer data further confirms that our simulated sleep–wake differences reproduce in vivo observations (Fig. 3d). In Fig. 3d, experimental measurements are shown with error bars, while simulation results include variability arising from parameter ranges. Despite this uncertainty, simulations remain within the experimental range and consistently predict higher electroosmotic velocities during sleep. Model predictions are robust to variations in key electrokinetic parameters, as discussed in the sensitivity analysis (Supplementary Fig. S2). These findings support the hypothesis that state-dependent neuronal activity directly modulates glymphatic flow via EO. ECS in the brain is not static but can undergo rapid structural changes, with its volume fraction expanding or contracting on the timescale of seconds during neuronal activity, seizures, or sleep–wake transitions ^41^. In the original analysis, we reported electroosmotic velocity (*v*) rather than volumetric flow rate (*Q*) to isolate the effect of electrokinetic forcing from geometric changes in the ECS. To further assess the impact of ECS expansion, we performed additional simulations with increased ECS volume fraction representative of sleep conditions. Consistent with electroosmotic theory, the results show that the average electroosmotic velocity remains unchanged (Supplementary Fig. S3), while the increased cross-sectional area leads to a proportional increase in volumetric flow rate (*Q* = *vA*). These results indicate that ECS expansion and electroosmotic driving act synergistically to enhance fluid transport during sleep, while the primary mechanism governing velocity remains the neuronal activity–dependent electrokinetic forcing. These results are summarized in Supplementary Table S2.

To assess the relative contribution of pressure-driven flow, we performed additional simulations incorporating a pressure gradient representative of arterial pulsation. We considered a gradient of 0.01 Pa/*µ*m, which is approximately two orders of magnitude larger than experimentally reported pressure gradients in brain parenchyma (∼0.1–0.3 Pa/mm) ^42^. Even under this elevated condition, the resulting interstitial velocities were on the order of 0.1 nm/s and remained approximately two orders of magnitude smaller than those generated by electroosmotic forcing (Supplementary Table S1). These results indicate that while pulsation may influence influx and modulate transport at the system level, pressure-driven flow alone is insufficient to drive significant bulk transport within the parenchymal ECS.

Beyond driving local parenchymal transport, EO intrinsically generates directional pressure gradients, which may critically contribute to coordinating other components of the glymphatic system. As illustrated in Fig. 4, CSF enters the parenchyma via periarterial spaces, traversing porous arterial boundaries before exchanging with interstitial fluid. Fluid must then exit through perivenous boundaries into perivenous spaces, ultimately joining venous efflux routes. Conservation of mass requires that these coupled compartments operate in balance, avoiding fluid buildup in any segment. Classical pressure-driven or diffusion-based models cannot readily explain how influx and efflux remain simultaneously regulated. Our results suggest that electroosmosis may provide a physical coupling between these processes, simultaneously supporting periarterial influx, parenchymal transport, and perivenous efflux through an activity dependent mechanism (Fig. 4).

**Figure 4:**
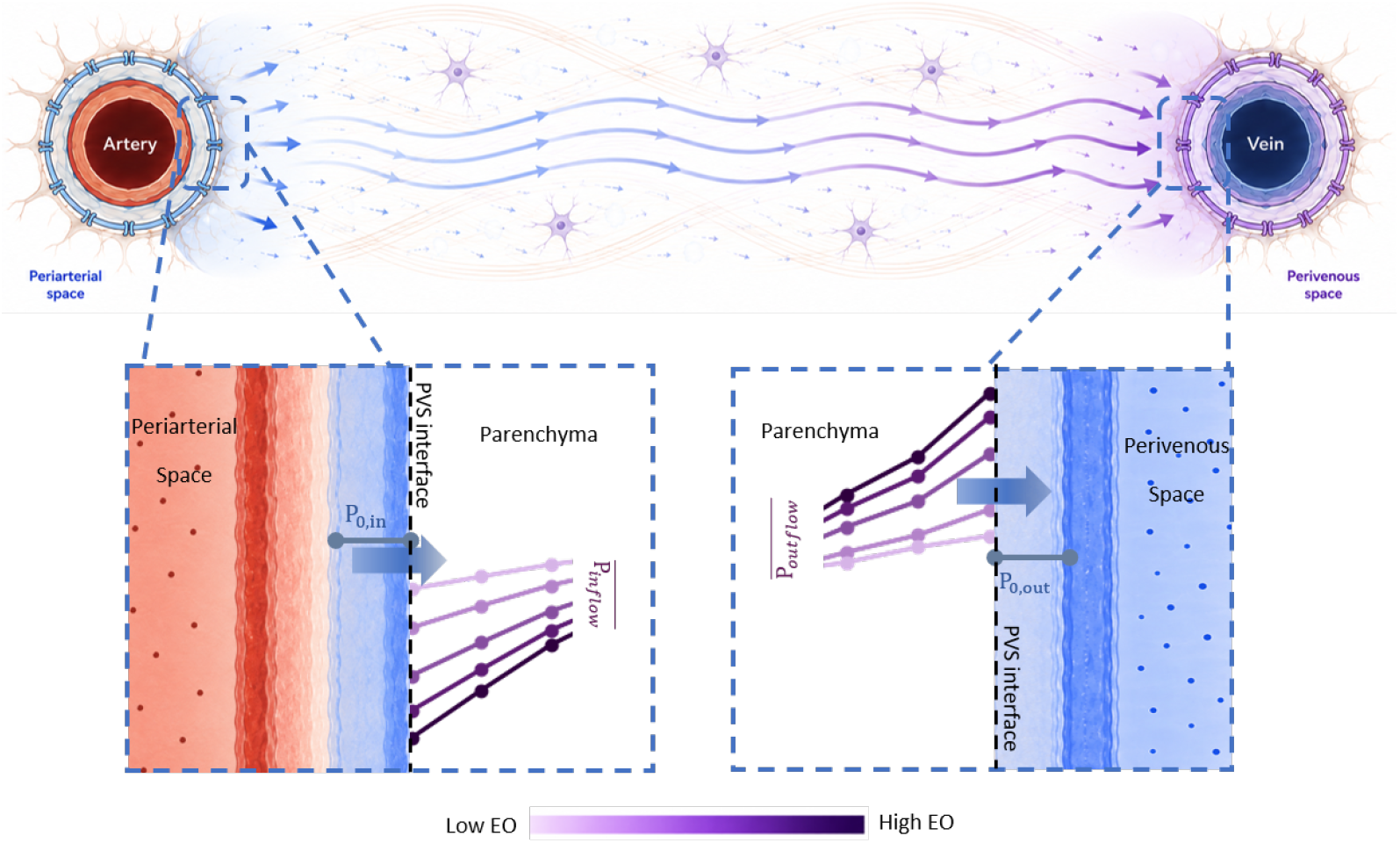
Electric field power regulates electroosmotic transport and boundary pressures. Schematic of glymphatic circulation showing CSF flowing from the periarterial space through parenchyma to the perivenous space. Top: streamlines illustrating transport from artery to vein across the parenchymal tissue. Bottom left: pressure profile at the periarterial (inflow) PVS interface showing a local pressure minimum (P0,in) that facilitates CSF entry; lines represent Low, Medium, and High EO conditions. Bottom right: pressure profile at the perivenous (outflow) PVS interface showing pressure buildup (P0,out) that scales with EO strength, facilitating CSF clearance.

Electroosmotic flow driven by locally generated electric fields induces spatial pressure variations within the tissue. These distributed pressure differences give rise to a relative pressure minimum near in-flow boundaries (e.g., periarterial regions) and a corresponding pressure elevation near outflow boundaries (e.g., perivenous regions). The pressure gradients emerge as a collective effect of spatially distributed electroosmotic forcing within the heterogeneous extracellular network. Neuronal activity propagates along axonal pathways and can generate locally directional extracellular currents; however, vascular and axonal orientations are not globally aligned, with vessels frequently traversing perpendicular to dominant fiber directions or varying along the same pathway ^43^. The local pressure minimum facilitates CSF entry through periarterial boundaries, while the elevated efflux pressure supports clearance through perivenous boundaries.

Thus, stronger neuronal activity, reflected in higher LFP wave power, not only enhances parenchymal flow but also amplifies periarterial influx and perivenous efflux through modulation of local boundary pressures. This provides a plausible mechanism for glymphatic efflux, a process that remains comparatively understudied and mechanistically unresolved in glymphatic physiology. Experimental studies have shown that chemogenetic inhibition of neuronal activity impairs glymphatic efflux and tracer clearance ^9^. To our knowledge, the present electroosmotic model is the first quantitative transport framework capable of explaining how neuronal activity can enhance parenchymal transport, and facilitate activity-dependent glymphatic efflux.

The pressure distribution shown in Fig. 4 has additional implications for solute clearance. As interstitial fluid moves from periarterial influx regions toward perivenous efflux pathways, metabolic byproducts and waste molecules are continuously collected and transported through the parenchyma. Consequently, solute concentrations are expected to increase along the transport pathway, leading to progressively larger osmotic pressures near downstream efflux regions. Such concentration gradients create a natural tendency for water to remain within the tissue and can therefore oppose fluid removal across efflux boundaries. The elevated pressures observed near the efflux regions in our simulations provide a potential means of overcoming this osmotic resistance while simultaneously facilitating exchange into downstream clearance pathways. In contrast, the reduced pressures observed near influx boundaries promote fluid entry from periarterial spaces. Together, these results suggest that the induced pressure field may serve a dual role: supporting hydraulic communication across compartment boundaries while maintaining effective clearance in the presence of solute concentration gradients that naturally arise during transport.

As shown in Fig. 4, pressure differences scaled with the power of applied LFP-derived fields, suggesting that neuronal oscillation intensity modulates not only parenchymal flow but also compartmental exchange across perivascular interfaces. This positions EO flow as an intermediate ‘pump’ that couples periarterial influx with perivenous efflux, amplifying glymphatic clearance during sleep when oscillatory power is greatest. We further considered the role of dynamic ECS volume, which expands and contracts across brain states ^41^. Because EO velocity depends primarily on surface electrokinetic forcing rather than absolute channel dimensions, our results remain valid, nonetheless, enhanced flow could itself promote ECS expansion, providing a feedback link between neural activity, electrokinetic transport, and microstructural remodeling.

Our findings establish electroosmosis as a viable and unifying mechanism for intraparenchymal interstitial fluid transport. By linking electric fields generated by neuronal activity to directional fluid motion in the ECS, our model explains key experimental observations, including enhanced glymphatic flow during sleep, modulation by brain state, and directional guidance by neuronal activation patterns ^9^. Supporting this connection, clinical monitoring of patients with TBI and subarachnoid hemorrhage has revealed that EEG activity can causally drive intracranial pressure fluctuations, indicating that neuronal electric fields directly modulate intracranial fluid dynamics ^44^. Unlike diffusion, or pressure-driven flow, electroosmosis uniquely provides directional, mass-conserving clearance that aligns with experimental data.

The brain is one of the most energy consuming organs of the human body, continuously generating metabolic waste. These include gases, metabolites such as lactic acid, misfolded or aggregated proteins, damaged organelles (e.g., mitochondria), and other macromolecules. These waste products are released into the interstitial space, where the glymphatic system facilitates their transport toward the venous circulation and ultimately into lymphatic pathways for clearance. The current understanding of glymphatic flow suggests that state dependent changes, including modulation of aquaporin-4 activity and ECS volume, contribute to enhanced fluid and waste transport, particularly during sleep. However, it remains unclear how waste is directionally transported through the highly tortuous parenchymal ECS. The mechanism proposed here provides this link: neural activity generates local extracellular electric fields which, in the presence of charged ECS interfaces, drive electroosmotic flow. This establishes a direct coupling between brain metabolism, through its regulation of neural activity, and fluid-mediated waste transport. Such a coupling may help explain how state dependent neural dynamics, including increased synchrony during sleep, contribute to enhanced clearance. It may also be particularly relevant in pathological conditions such as traumatic brain injury and neurodegenerative disease, where both electrical activity and fluid transport are disrupted. Furthermore, recent reports demonstrating that focused ultrasound can enhance glymphatic flow may be consistent with modulation of electrokinetic mechanisms, providing additional context for the relevance of the proposed framework.

In parallel, accumulating experimental evidence indicates that brain fluid transport can be actively modulated by physiological and external factors. Mechanical stimulation such as ultrasound has been shown to enhance glymphatic transport and interstitial tracer movement^45^, while modulation of vascular pulsation can accelerate interstitial fluid drainage ^46^. More recently, electrical neuromodulation approaches such as transcranial direct current stimulation have been reported to alter CSF–ISF exchange in vivo ^47^. These findings collectively suggest that brain fluid transport may be influenced by multiple dynamic driving mechanisms beyond passive diffusion alone. The electroosmotic mechanism proposed here represents a complementary biophysical pathway that could operate within the charged extracellular microenvironment to couple neuronal electrical activity with interstitial fluid transport. At a broader level, mechanisms that enhance clearance efficiency, such as electroosmosis, may have influenced the evolutionary optimization of brain morphology and glymphatic circulation^48^. Beyond resolving a longstanding gap in glymphatic physiology, this insight opens translational opportunities. For example, weak transcranial alternating current stimulation at slow-wave frequencies could enhance electroosmotic transport, accelerating clearance during sleep or aiding drug delivery ^35^. Such approaches would leverage the brain’s endogenous mechanism rather than imposing external pressure or diffusion, offering a novel therapeutic pathway for neurodegenerative disease. Future work could investigate whether propagating neuronal activity patterns, such as those observed in epilepsy, give rise to directional interstitial fluid transport. The proposed electroosmotic mechanism suggests this as a testable hypothesis that can be examined using combined measurements of neural activity and fluid flow. The ROM framework developed here provides a practical path toward this goal, enabling tissue- and organ-scale simulations that would otherwise be computationally intractable.

Our findings support a conceptual separation between perivascular circulation and intraparenchymal transport. Existing experimental evidence strongly suggests that arterial pulsations, vasomotion, and related vascular dynamics contribute to fluid movement within periarterial and perivenous spaces. However, the parenchymal extracellular space represents a distinct transport environment characterized by nanoscale confinement, high tortuosity, and substantially greater hydraulic resistance. In this context, intraparenchymal transport is unlikely to be a simple continuation of flow established within the perivascular compartments. Rather, perivascular circulation may serve primarily to deliver and collect fluid at tissue boundaries, while a separate transport mechanism governs movement through the extra-cellular space itself. This distinction helps reconcile how robust circulation can occur within perivascular pathways while pressure-driven transport remains insufficient to account for the observed rates of interstitial fluid movement within the parenchyma. Viewed in this framework, perivascular and parenchymal transport are complementary processes operating in hydraulically distinct compartments that together contribute to whole-brain glymphatic circulation.

## Methods

### Geometry Generation

To reconstruct a realistic 3D geometry of the extracellular space, we used publicly available anatomical data reported by Kinney et al. ^32^, who reconstructed hippocampal neuropil from serial electron microscopy images. We downloaded the co-registered EM image stack, segmentation contours, and 3D surface models provided. The original EM images were traced using RECONSTRUCT, and 3D surfaces were generated with VolumeRover, as detailed in the original publication.

We began with the STL surface mesh of the ECS-accessible space extracted from the provided data. Due to the highly complex, high-resolution topology of the mesh, it was not immediately suitable for computational fluid dynamics simulations. To prepare the geometry, we first imported the mesh into ParaView to crop a representative 4 *µ*m^3^ volume that preserved key structural features while reducing computational burden.

The cropped geometry was then imported into Blender for extensive cleanup. Each cellular component was isolated and inspected to correct inverted normals, fill holes, and eliminate surface artifacts. We performed localized remeshing and smoothing to improve watertightness and reduce mesh irregularities. Because our simulations targeted fluid transport, we applied a Boolean inversion to extract the fluid-accessible domain (i.e., the void space between cells).

This inverted, cleaned fluid domain was imported into SolidWorks, where we repaired non-manifold edges and ensured geometric validity for mesh generation and converted to a solid part. The final solid model was imported into COMSOL Multiphysics, where tetrahedral meshing with local refinement was used to resolve narrow ECS channels and enable stable electroosmotic simulations.

This reconstruction pipeline—from EM image-based anatomical data to simulation-ready geometry—enabled us to model flow through a physiologically grounded ECS microenvironment. Code for geometry handling and STL processing is available upon request.

### Simulation Setup

The fluid motion was modeled by the incompressible Navier–Stokes equations in three dimensions:

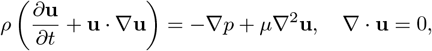

where **u** is the velocity field, *p* is pressure, *ρ* is the fluid density, and *µ* is the dynamic viscosity. This formulation directly solves for the solvent velocity and pressure fields in the extracellular space. Electroosmotic effects are incorporated through a slip velocity boundary condition, which represents the net action of electric forces on ions within the thin electric double layer and their viscous coupling to the bulk fluid. Due to the microscale geometry of the extracellular space and low flow velocities, the Reynolds number remained small throughout, allowing us to neglect inertial effects. This reduces the system to the unsteady Stokes equations:

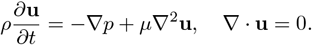

Electroosmotic flow was modeled through a wall slip velocity condition. Specifically, the Helmholtz–Smoluchowski formulation was applied on all solid walls in the ECS geometry:

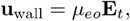

where 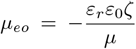 is the electroosmotic mobility, with *ε*_*r*_ and *ε*_0_ being the relative and vacuum permittivity of the fluid, *ζ* the wall zeta potential, and *µ* the viscosity. The tangential electric field is defined as

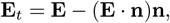

where **E** is the applied electric field and **n** is the unit normal to the surface.

The electric field E(t) was modeled as spatially uniform within each computational domain but varying in time. The temporal waveform was derived from experimentally measured extracellular field potentials (local field potentials, LFPs) reported by Jiang-Xie et al. ^9^, obtained using ultrathin electrodes positioned within the brain extracellular space. These measurements represent extracellular electric fields acting on ions in the interstitial fluid, rather than intracellular membrane potentials. Because the computational domains considered here are on the order of a few micrometers, which is significantly smaller than the spatial scale over which LFPs vary (∼ 100–200 *µ*m), the electric field can be reasonably approximated as locally uniform within each domain. This scale separation is consistent with standard electrokinetic modeling in porous and confined systems, where microscale transport is evaluated under locally applied fields. Under this framework, electroosmotic velocity scales linearly with the applied electric field via the Helmholtz–Smoluchowski relation. Therefore, the use of experimentally measured extracellular field amplitudes provides a physiologically grounded estimate of the electroosmotic driving force. The time-dependent waveform was varied in both shape and amplitude to examine its effect on electroosmotic pressure gradients and net transport, as described in subsequent sections.

The simulations were performed in COMSOL Multiphysics using the Laminar Flow interface with a time-dependent solver and variable step size to resolve transient dynamics accurately. To validate the numerical implementation of electroosmotic flow, we reproduced a canonical pressure-free electroosmotic flow in a charged microchannel and compared the simulation results with experimental measurements reported in the literature. The model accurately recovers the expected velocity profiles and agrees well with experimental data ^38^ (see Supplementary Information, Fig. S1), confirming the validity of the electrokinetic formulation used in this study.

### Mesh Convergence Analysis

To ensure the accuracy and stability of the numerical simulations, a mesh refinement study was conducted on the extracellular space geometry. Multiple unstructured tetrahedral meshes were generated with progressively smaller average element sizes. Key output metrics were evaluated at each refinement level. Convergence was assessed by comparing these metrics across mesh resolutions. The simulation results were shown mesh-independent when successive refinements produced changes below a predefined threshold. Final simulations were performed using the coarsest mesh that satisfied this convergence criterion to reduce computational cost while maintaining accuracy.

The final mesh used in all simulations consisted of 2,380,925 vertices and 8,948,873 elements (7,127,270 tetrahedra, 13,614 pyramids, 180,798 prisms, and 107,012 triangles)

### Domain Size Sensitivity

Because the extracellular space structure was sampled from a finite subvolume, it was necessary to evaluate whether the simulated flow behavior was sensitive to the domain size. To assess this, simulations were performed on a geometry with twice volume while keeping the applied electric field and boundary conditions fixed.

Flow metrics were compared across different domain sizes to determine whether the results were size-independent. The selected domain for final simulations was chosen to be the smallest size for which increasing volume no longer significantly affected the outcome.

### Electric Field Power and Waveform Generation

To characterize how electrophysiologically driven electric fields influence electroosmotic flow, we generated synthetic, time-varying electric field signals based on physiological LFP recordings. Experimental LFP data from rodent cortex during sleep and wake states were analyzed in the frequency domain using Fourier transforms to determine the typical spectral content and power distribution in each state.

Rather than applying a single frequency or fixed waveform, we constructed a family of time-varying electric fields with varying combinations of amplitude and frequency. All generated signals were sinusoidal in shape but differed in these parameters to span a continuous range of field powers., encapsulating contributions from both field amplitude and frequency when considered across a waveform family.

These synthetic fields were applied as time-dependent inputs via the electroosmotic slip boundary condition. The resulting EO response, quantified by pressure gradients and flow rates, was then evaluated as a function of the driving field power.

### ROM Development

Microstructure resolved simulations, while mechanistically accurate, are computationally limited at tissue scales due to the complexity and resolution of the ECS geometry. A reduced order model addresses this by replacing the fully resolved geometry with a computationally efficient surrogate that captures the essential physics. Instead of resolving individual pore-scale features, a ROM operates on a homogenized continuum representation of the domain, parameterized by effective properties extracted from the full model. This makes ROM simulations orders of magnitude faster than their microstructure-resolved counterparts, enabling tissue-scale predictions that would otherwise be the. The key requirement for a valid ROM is that it reproduces the outputs of the high-fidelity model across the full range of physically relevant conditions, not just at a single operating point, ensuring the reduced representation captures the underlying physics.

To extend electroosmotic transport predictions to tissue scales, a physics-informed, data-driven reduced-order model was developed. The model combines a continuum porous medium representation of the ECS, whose governing equation structure is derived from physical principles, with an effective transport coefficient computed from microstructure-resolved simulation data.

The anatomically reconstructed ECS geometry was replaced by a homogenized continuum block of equivalent volume. The porous medium was characterized by porosity *ε*_*p*_ = 0.317 and permeability *κ* = 9.7 × 10^−17^ m^2^, Fluid flow was governed by the Brinkman equations:

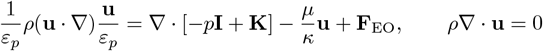

where 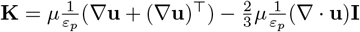 is the viscous stress tensor.

Electroosmotic transport was incorporated as a volumetric body force applied throughout the porous domain:

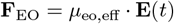

where **E**(t) is the time-varying electric field derived from experimental LFP recordings, and *µ*_eo,eff_ is an effective electroosmotic body force coefficient with units of N m^−3^ per V m^−1^. This coefficient encapsulates the combined influence of pore geometry, surface charge distribution, and tortuosity.

The value of *µ*_eo,eff_ was computed from the microstructure-resolved simulation data by finding the value that minimized the discrepancy between ROM-predicted volume-averaged velocities and exact simulation outputs under identical electric field inputs. A single value of *µ*_eo,eff_ = 19.88 N m^−3^ V^−1^ m was found to reproduce exact simulation results across the full range of tested electric field magnitudes and brain-state waveforms (Fig. 2e). The parameter was applied without modification to all tissue-scale ROM simulations.

## Acknowledgments

The authors would like to thank the UCLA Brain Injury Research Center, the UCLA Steve Tisch BrainSPORT Program, and the UCLA Easton Brain Health Program for their support and resources. P.H. acknowledges support from the Translational Neurotechnology (TNT) program through a T32 NIH Fellowship. The authors are grateful to Dr. Jeffrey W. Ruberti, Dr. John Hollister, and Dr. John K. Lee for their fruitful discussions and valuable suggestions that greatly strengthened this work.

## Availability of data

The datasets used and analysed during the current study are available from the corresponding author on reasonable request.

## Author Contributions

P.H. and H.P.K. contributed to conceptualization and methodology. P.H. performed simulations, validation, data analysis, and visualization. H.P.K. contributed to simulation design, provided resources, intellectual guidance, and supervised the study. P.H. and H.P.K. wrote the manuscript with input from all authors. A.C.W., M.L.P., and C.C.G. contributed clinical and physiological expertise and intellectual guidance.

