## Supplementary material for "Neuronal Activity-Dependent Electroosmosis and Its Potential Role in Interstitial Fluid Flow in Glymphatic System": Supplemenatry Figures and Table

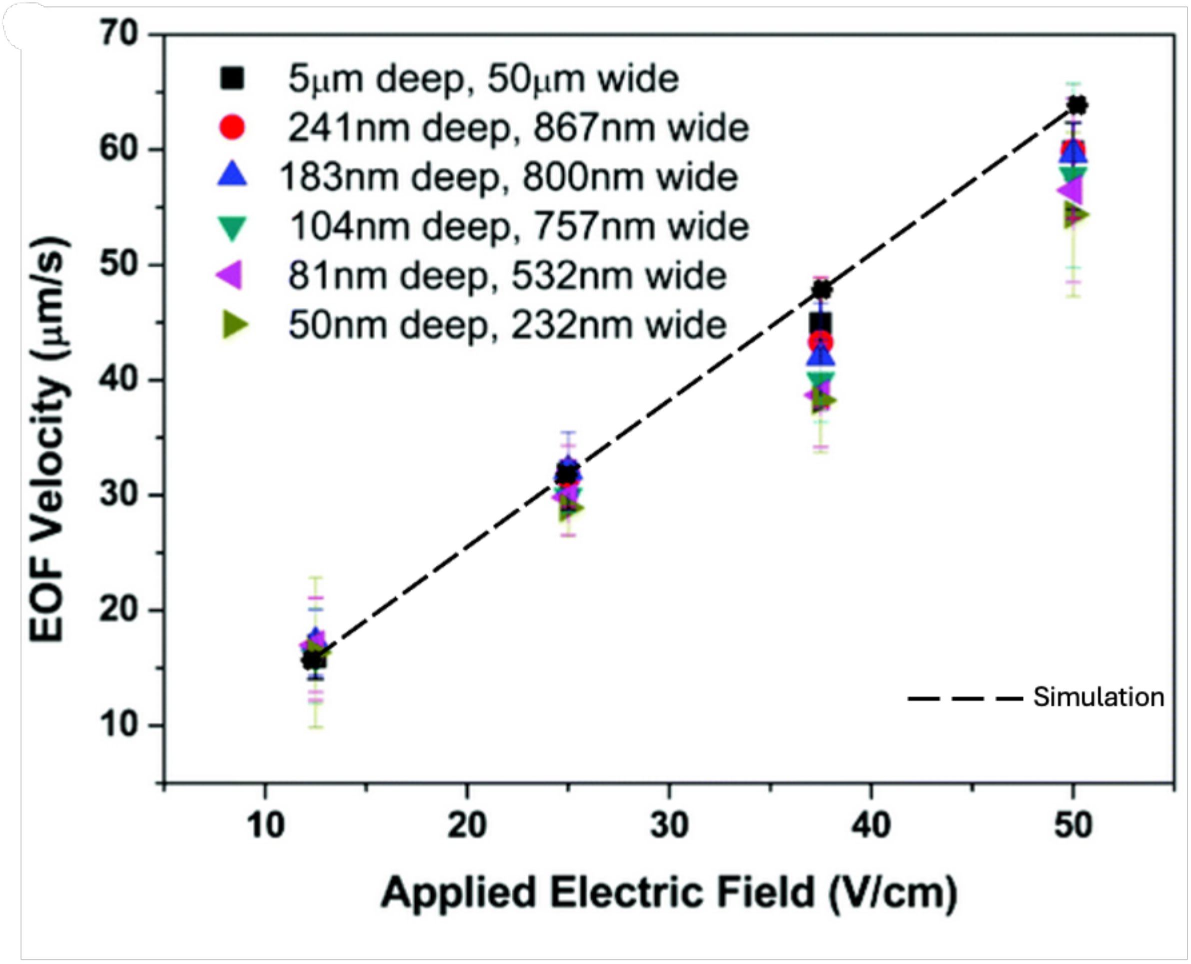


**Fig. S1.** Validation of the electroosmotic model against experimental measurements reported by Peng et al. Electroosmotic velocities are shown as a function of applied electric field for channels spanning micro- to nanoscale dimensions (as indicated in the legend). Symbols represent experimental data, while the black dashed line denotes simulation results based on the present model. The close agreement across channel sizes supports the applicability of the electroosmotic formulation in confined geometries relevant to the extracellular space.

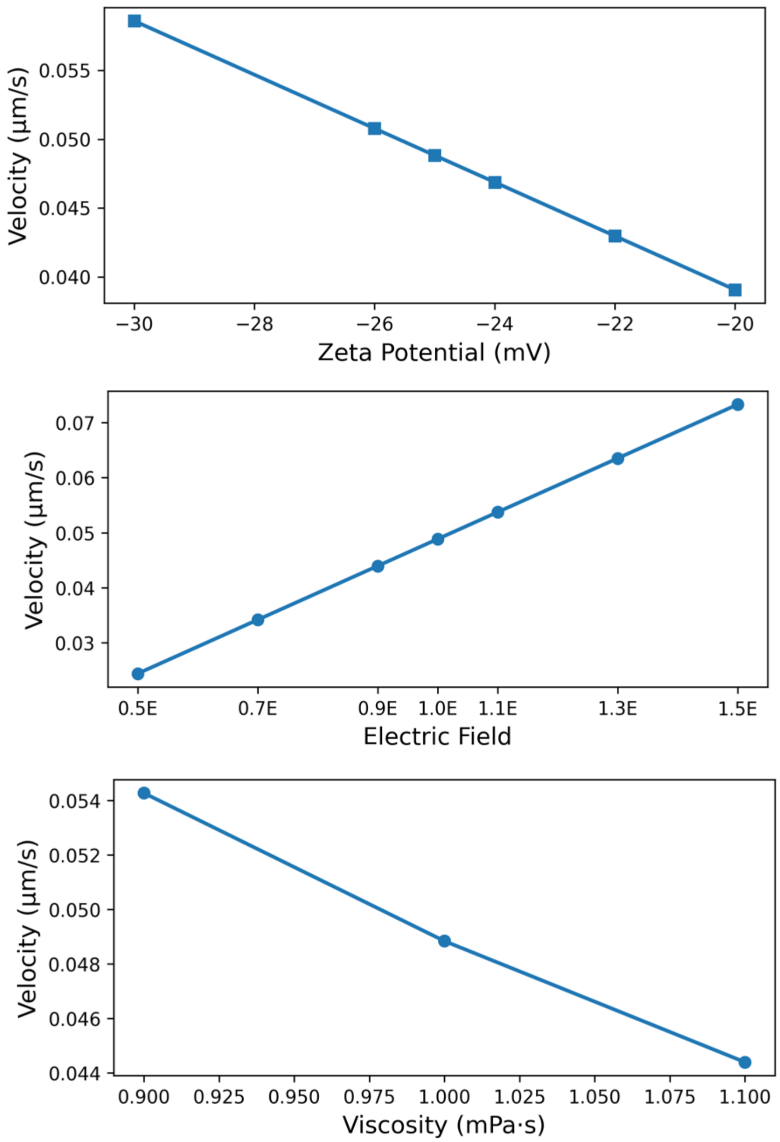


**Fig. S2.** Sensitivity analysis of electroosmotic velocity in the 3D simulation model. The dependence of velocity on key electrokinetic parameters is shown: (top) zeta potential, (middle) applied electric field, and (bottom) fluid viscosity. The results demonstrate the expected electroosmotic scaling, with velocity increasing linearly with electric field, decreasing with increasing viscosity, and varying proportionally with zeta potential, consistent with the Helmholtz–Smoluchowski relation, confirming that the numerical implementation reproduces the expected electroosmotic behavior.

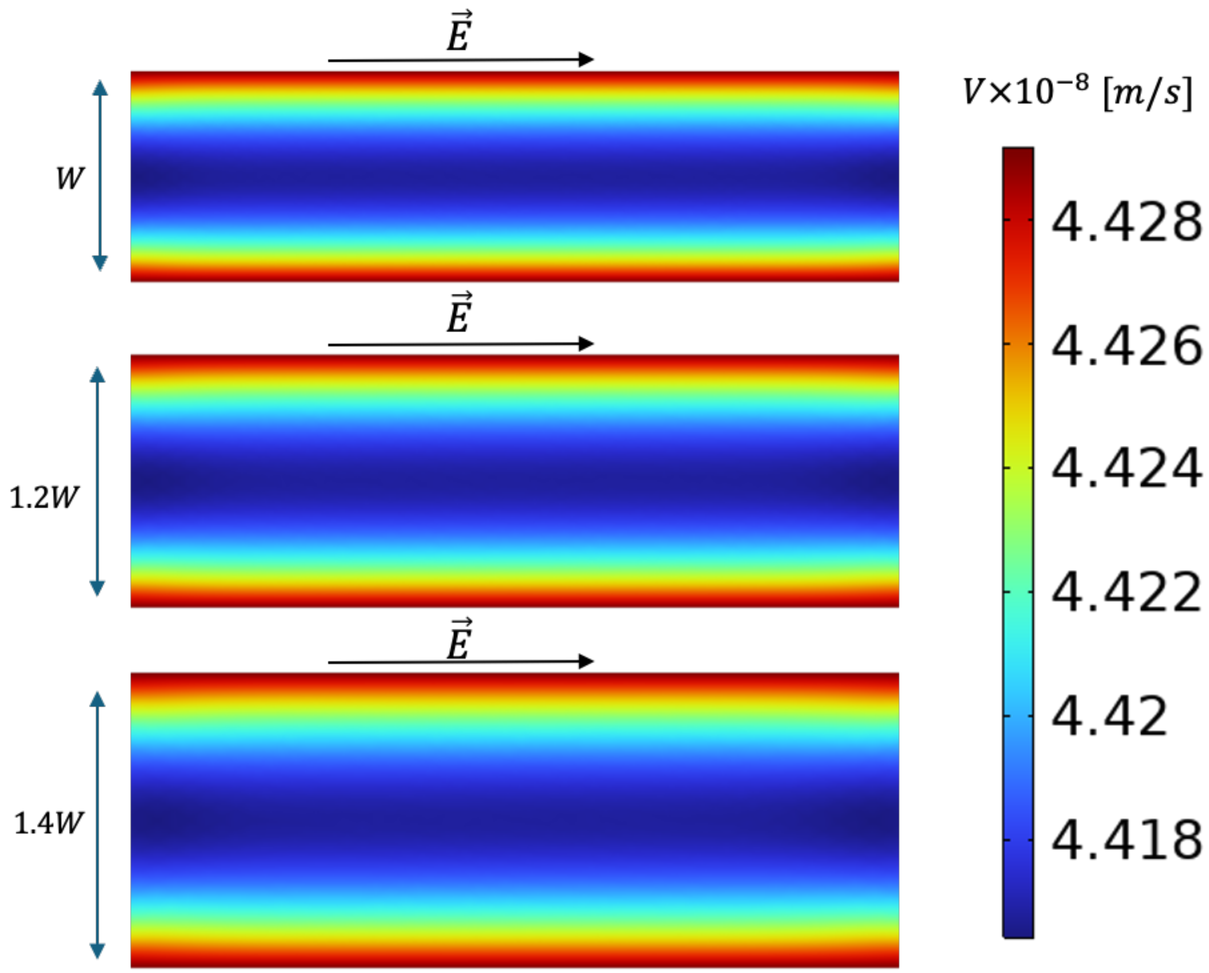


**Fig. S3.** Independence of electroosmotic velocity on channel width at spatial and electric field scales relevant to ECS . Velocity magnitude is shown for channels of increasing width (indicated by arrows) under identical electrokinetic conditions. Despite changes in channel size, the velocity profiles remain consistent, demonstrating that electroosmotic velocity is independent of channel dimensions, as predicted by the Helmholtz–Smoluchowski relation.

**Table S1.** Model robustness under different boundary conditions. Average electroosmotic velocity within the domain is reported for a range of boundary conditions, including inlet flow, open, periodic, and pressure-driven configurations. The results show consistent velocities across boundary conditions when electroosmotic forcing is present, while pressure-driven flow alone produces velocities several orders of magnitude smaller.

| Boundary Condition | Average Velocity (m/s) |
|---|---|
| Inlet flow and outlet pressure | $4.8 \times 10^{-8}$ |
| Open Boundary | $4.57 \times 10^{-8}$ |
| Pressure Gradient + EO | $4.62 \times 10^{-8}$ |
| Periodic boundary | $7.41 \times 10^{-8}$ |
| Pressure Driven Only, no EO | $8.74 \times 10^{-10}$ |

**Table S2.** Effect of extracellular space expansion on electroosmotic velocity. Average velocity within the domain is reported for increasing ECS expansion factors. The results show that electroosmotic velocity remains unchanged with increasing ECS size, consistent with theoretical predictions, while implying an increase in volumetric flow rate due to the larger cross-sectional area.

| ECS Expansion Factor | Average Velocity (m/s) |
|---|---|
| 1.0 | $4.8 \times 10^{-8}$ |
| 1.2 | $4.8 \times 10^{-8}$ |
| 1.4 | $4.8 \times 10^{-8}$ |
